# Many facets of self-directed behaviours in macaques

**DOI:** 10.1101/2025.08.11.669676

**Authors:** Debottam Bhattacharjee, Eva S.J. van Dijk, Roos E.M. de Vries, Charlotte E. Kluiver, Mia Valomy, Adam N. Zeeman, Jorg J.M. Massen

## Abstract

Emotional states like anxiety are adaptive, assisting individuals during social uncertainty and threats. While humans can self-report anxiety, assessing comparable states in animals requires objective measures. Self-directed behaviours (SDBs), including self-scratch, autogroom, and yawn, are indicators of anxiety, particularly in nonhuman primates. Yet it is unclear whether they reflect a common underlying process or distinct responses shaped by demography, social interactions and status, and the broader social dynamics. We studied 13 macaque groups across six species along a *despotic–egalitarian* social style gradient to investigate the factors associated with variation in SDBs. Our findings revealed partial correlational consistency among the SDBs, where only self-scratch and autogroom positively correlated. Further, we found that males yawned more than females, and individuals receiving less aggression exhibited longer autogroom bouts. Dominance rank showed behaviour-specific effects, with weak evidence for an association with self-scratch, longer autogrooming in subordinates, and a non-linear pattern in yawning, which was lowest at intermediate ranks. Alpha status was linked to more frequent yawning than others. And finally, social style showed no consistent effects on SDBs. Our research demonstrates that SDBs in macaques reflect differential responses related to social uncertainty rather than one anxiety dimension, providing insights into primate emotion regulation.

## 1. Introduction

Individual emotional states are mediated by physiological and behavioural changes across species, offering benefits that range from welfare to survival [1,2]. In social contexts, a common emotional state is anxiety – an individual’s response to aversive situations or threats [3,4]. While adult humans can self-report their experience of anxiety, assessing this psychosocial or affective state in animals requires objective methodological approaches. Self-directed behaviours (SDBs), defined by behaviours directed toward the self, such as self-scratch, autogroom, and yawn [5], are powerful and widely used proxies to capture anxiety-related emotional states in animals [6]. Although these behaviours can serve varying functions, from environmental to hygienic, their social functions have received considerable attention [6–10]. By testing the discriminant validity of SDBs, previous research has found their positive correlations with physiological measures of anxiety and broadly ‘stress’. For instance, post-conflict SDBs positively correlated with maximum heart rate in greylag geese (*Anser anser*) [11]. In European starlings (*Sturnus vulgaris*), acute crowding led to elevated SDBs, heart rates, and corticosterone concentrations [12]. SDBs increased when long-tailed macaques (*Macaca fascicularis*) were given anxiety-inducing drugs [13,14]. However, a lack of positive correlation between SDBs and those physiological measures was also reported (see [15–17]). Nonetheless, research on various social contexts, such as during allogrooming and conflict, has found systematic fluctuations in SDBs [18–21], establishing their mediating roles in anxiety. Thus, the consensus is that SDBs are displacement behaviours that do not necessarily reflect chronic stress (but see [22]) and are instead associated with transient or short-term emotional states with *coping* benefits [5,6,11,17,23–27], for example, helping individuals overcome challenges related to stressful situations (e.g., during intragroup conflicts). Yet, considerable between-individual variation in the expression of SDBs is prevalent, and comprehensive evidence about their underlying proximate mechanisms is still poorly understood.

Demographic factors such as age and sex, as well as social factors like aggression, are known to be associated with SDBs and their variations. A decline in physical conditions with ageing can be associated with higher negative affective states [28]. Consequently, older individuals may use SDBs as a functional means of coping, or they may increase reliance on frequent hygienic activities [8,29]. By contrast, general life experience with ageing may lead to better emotional regulation [30], where individuals become more certain about their relationships with group members, potentially leading to a decrease in SDBs. Sex-level differences have also been reported, with males often exhibiting more SDBs than females [31]. For instance, yawning in males can lead to the exposure of large canines, which carry social information on dominance and promote group stability. Inter-sex competition, including aggression [32], can also potentially lead to variations in SDBs. Thus, sex-level differences can be explained by anatomical (*sensu* sexual dimorphism), physiological (such as concentration of androgens), as well as social (such as competition and aggression) factors [32–34]. However, sex-based variations in SDBs are not conclusive, as comparable rates between females and males have also been reported (see [35,36]). Moreover, aggression can influence SDBs, irrespective of sex differences [8,37–39], with a positive association between receiving frequent aggression and exhibiting higher rates of SDBs. Furthermore, many studies have analysed SDBs as a single composite index (e.g., see [17,29,40,41]), based on the assumption that the elemental behaviours (i.e., self-scratch, autogroom, and yawn) covary, or simply due to limited sampling effort, resulting in insufficient data on each behaviour separately. Overall, while there is still a critical knowledge gap regarding how age, sex, and aggression are associated with specific elemental anxiety-related behaviours, the existing inconclusive evidence also points to the need to investigate other social factors.

In stable social hierarchies, social status or dominance rank may influence SDBs. In particular, the broad mechanisms – (un-)certainty in the outcomes of within-group social interactions and rank-related energetic constraints – can be grouped under two non-mutually exclusive hypotheses. The uncertainty hypothesis suggests that subordinate group members experience greater unpredictability in the outcomes of social interactions and therefore express more frequent SDBs, as these behaviours help them cope with tense situations or avoid conflict [38,41–45]. Particularly, such ‘reactive’ coping benefits may come from heightened ‘flight’ responses, helping subordinates to avoid higher-ranking group members and potential conflicts, or to seek support from closely affiliated group members [6,38,46–48]. By contrast, the energetic constraint hypothesis posits that different ranks within a stable hierarchy are associated with unique costs and benefits, and that stress-related responses covary with rank [49]. Accordingly, the top rank or the ‘alpha status’ is linked to elevated energetic demands (*sensu* ‘executive stress syndrome’) [49], from physiological costs of mating and functional needs for vigilance (e.g., against outgroup threats, but also intragroup competitors/challengers) to group maintenance. Empirical findings across primates reflect both mechanisms, leading to inconclusive patterns: in some systems, high-ranking individuals are the most stressed, particularly when dominance is maintained through frequent physical aggression; in others, lower-ranking individuals show more stress when hierarchies rely primarily on intimidation; and in more egalitarian societies, rank-related stress differences may be weak or absent [50]. These contrasts highlight that the relationship between rank and stress may depend on the species’ social style, the stability of dominance relationships, and the broader dynamics of within-group interactions [50,51]. However, comparative research investigating SDBs across phylogenetically close species with varying social styles is limited.

The genus *Macaca* represents ∼22 species of group-living primates, with similar social organizations yet strikingly different social tolerance levels or social styles [52–54], providing an excellent opportunity to investigate SDBs and their proximate mechanisms comparatively. The broader social styles (interchangeably referred to as tolerance grades) in macaques are shaped by phylogenetically co-varying structural traits [52–55], primarily reflected in how individuals tolerate one another within a group. These co-varying traits include hierarchical steepness of dominance, coalitionary tendencies, and nepotistic and kin biases. While robust comparative empirical evidence is still scarce, some studies indicate that these traits indeed covary, and macaque species or societies can be categorized into four distinct tolerance grades along a despotic–egalitarian gradient [56–58]. Despotic societies are primarily characterized by steeper dominance hierarchies and more frequent aggressive interactions than egalitarian ones [56,59]. From grade one (or the despotic end) to four (or the egalitarian end), social tolerance is considered to increase (but see [60]). Accordingly, primatologists use these grades to distinguish the different macaque species and to test various ecological, social, and cognitive hypotheses. As varying social styles can influence individuals at the physiological [61], behavioural [62], and cognitive [63] levels, SDBs can also be potentially shaped by them, and by studying SDBs at the grade level, we can account for the phylogenetic inertia. Furthermore, rank-related uncertainties are higher in more egalitarian than in despotic societies, which are often described as “complex” with regard to within-group communication and interaction [64,65]. Yet, whether SDBs generally vary across social styles remains a rarely investigated topic. Therefore, a comparative assessment of SDBs across macaque societies representing the four social styles can provide critical insights into their functions and, broadly, evolution.

Here we comparatively investigated three different SDBs (self-scratch, autogroom, and yawn) across six macaque species: Japanese (*Macaca fuscata*), rhesus (*M. mulatta*), long-tailed (*M. fascicularis*), lion-tailed (*M. silenus*), barbary (*M. sylvanus*), and crested (*M. nigra*) macaques. These species represented all four social styles (Grade 1: *M. fuscata* and *M. mulatta*; Grade 2: *M. fascicularis*; Grade 3: *M. silenus* and *M. sylvanus*; Grade 4: *M. nigra*, cf. [52–54]). Using a comprehensive dataset including 113 individuals (>4 years of age), we investigated (i) whether the three SDBs covary or exhibit correlational consistency, and (ii) the associations of SDBs with age, sex, aggression, dominance rank, and social styles or grades. Although we used SDB to denote the three behaviours, we analysed them separately and at the behaviour level.

We made a two-tailed prediction for age: older individuals would exhibit either higher (due to the general decline of physical conditions leading to increased hygienic requirements) or lower (due to better emotion regulation and more certainty of social relationships) amounts of SDBs. Aligning with the dimorphism hypothesis, we expected that SDBs, particularly yawn behaviour, would differ between females and males, with males exhibiting higher SDBs than females [33,34]. We expected a relationship between aggression and SDBs, specifically with individuals receiving more frequent aggression exhibiting higher amounts of SDBs. Due to the non-mutually exclusive nature of the uncertainty and energetic constraint hypotheses, we expected complex non-linear relationships between ranks and SDBs (cf. [49]). On one hand, we predicted that lower-ranking individuals would display higher amounts of SDBs than their higher-ranking group members due to uncertainty in the outcomes of social interactions. On the other hand, if higher ranks are associated with unique costs, such as elevated energy demands for mating and vigilance, and/or exhibition of dominance (cf. [49]), then the highest ranks or the alpha status would be associated with higher amounts of SDBs than the rest of the group members. Finally, for social styles, we had two alternative predictions. If despotism intensifies social stress through frequent aggressive interactions and intimidation (cf. [52,59]), then individuals in more despotic societies are expected to exhibit higher rates of SDBs than those in more egalitarian societies. Alternatively, if egalitarian societies are characterized by more fluid and less predictable (i.e., more complex) social relationships (cf. [41,63]), individuals in these societies would experience higher social uncertainty than those in despotic societies, subsequently leading to increased SDBs.

## 2. Materials and Methods

### 2.1. Study subjects

We observed 13 captive macaque groups across six species—*Macaca fuscata*, *M. mulatta*, *M. fascicularis*, *M. silenus*, *M. sylvanus*, and *M. nigra*—in their existing social environments. See **Table S1** for details on the subjects, and housing and husbandry protocols. Only individuals ≥ 4 years – considered sexually mature [66,67] – at the time of testing (n = 113) were included in the study and analyses (see **Table S1**). All groups had unrestricted access to fresh drinking water. No alterations were made to feeding protocols for this research. The animals were well habituated to the presence of the observers and exhibited no signs of stress during data collection.

### 2.2. Behavioural data collection using continuous focal sampling

Behavioural observations were carried out using a continuous focal sampling method between late 2020 and early 2023. We conducted these observations from 0800 to 1830 hours. Each focal observation lasted 20 minutes without periods when the individual was out of sight. The order in which focal animals were observed was randomized. From the 113 individuals (≥ 4 years, range: 4 – 29 years; female = 92, male = 21), we obtained a total of 31,480 minutes of behavioural data (**Table S1**). The mean (± SD) observation minutes per individual was 278.58 ± 75.23 (range = 160 – 460 min). The number of observation minutes differed across groups, largely due to the varying group sizes. Focal data were used to assess SDBs, aggressive behaviours, and dominance ranks.

### 2.3. Self-directed behaviours

We investigated three widely recognised SDBs – self-scratch, autogroom, and yawn [5,6,13,16,23,41,68]. Self-scratch was defined as the act of an individual using their finger, hand, or foot to rake across their own skin. Autogroom was defined by an individual grooming themselves by closely inspecting the skin, hands, nails, toes, or other body parts. Finally, we employed the widely accepted definition of a yawn (cf. [68]) – an involuntary and powerful gaping of the jaw accompanied by deep inspiration, followed by a temporary period of peak muscle contraction and passive closure of the jaw with shorter expiration. We could not differentiate between self-directed and directed display yawns, as they are challenging to disentangle in social contexts [69]. We calculated the rates of self-scratch and yawn (frequency per minute) separately to control for the varying observation minutes across groups and species. For autogroom, we quantified the duration and controlled for the observation variation by calculating seconds per minute. A new autogroom bout was considered if it was resumed after an interval of at least 5 seconds to account for data independence.

### 2.4. Aggression initiated and received

We extracted data on five behavioural variables (*lunge, open-mouth threat, physical attack/hit*, *bite,* and *chase*, see [70] for definitions) that represented both contact– and non-contact aggression. For each individual, we calculated the rates (frequency per minute) of aggression initiated (toward others) and aggression received (from others). By calculating rates, we controlled for the varying observation minutes across groups and species.

### 2.5. Dominance rank and social style

Based on five variables that represent unprovoked submissive behaviours (*avoid*, *be displaced*, *silent-bared teeth*, *flee*, and *social presence*, see [70] for definitions), we assessed individual dominance (or social) ranks as well as the steepness of the within-group dominance hierarchies using a Bayesian Elo-rating method. For all groups, except the *M. fuscata*, we used previously published data to obtain these values [60]. Individual dominance ranks were obtained as ordinal ranks at the group level. For *M. fuscata*, the limited number of dyadic unprovoked submissive interactions could not be used to calculate the ordinal ranks reliably. Therefore, we used previously published results on the matrilines to which those individuals belong [71]. Since the sampled individuals were adult females only in the *M. fuscata* group (cf. **Table S1**), matrilineal rank inheritance can be assumed [72], and these alternative measures can be useful for determining the ordinal ranks of the individuals. Nonetheless, we took caution while performing statistical analyses using this group in some cases. Due to the relatively long-term data collection (spanning ∼8 months per group), hierarchies were assumed to be stable, and no rank reversals were observed during the study.

The steepness of hierarchies followed the assumption of the co-variation framework (cf. [52–54]) that more despotic societies show higher steepness than more egalitarian ones [60]. Previously published data on our study groups revealed a moderate to strong correlation between macaque tolerance grades (or social styles) and hierarchy steepness (with all individuals: *r* = –0.33; with only adult females: *r* = –0.61) [60]. The fulfillment of this assumption, based on empirical data, validated our assignment of social styles to the species we investigated.

### 2.6. Data coding

Video-recorded data were coded using BORIS (version 7.13.6) or in a frame-by-frame manner using PotPlayer (version 240618). Due to data coding by multiple trained coders (by D.B., E.S.J.vD., R.E.M.dV., C.E.K., M.V., and A.N.Z., with assistance from 14 more coders), we comprehensively checked for inter-rater reliability. The agreement check was performed after all the coders became fully trained to code behavioural data. Overall, the agreement was high (Intraclass correlation coefficient or ICC(3,k) = 0.88, including SDBs, dominance rank, and aggression). Particularly, for the SDBs, the ICC(3,k) value was 0.94, suggesting excellent agreement among all coders.

### 2.7. Statistical models and packages

All statistical analyses were conducted in R (version 4.1.1) [73]. We used a Bayesian modeling approach using the “*brms*” package [74]. SDBs (response variables) were modelled using multivariate Gaussian models with identity links, which allowed them to be estimated jointly with a residual correlation structure. Three models were constructed to test the predictions of the study. In all models, we included a nested random effect of species and group (i.e., group within species) to account for the hierarchical sampling structure.

*Model 1* was used to investigate the association between the three SDBs, specifically whether they exhibited strong correlational consistency. We used the default weakly informative priors implemented in *brms*: flat priors for intercepts, half-student-t (3, 0, 10) priors on the residual standard deviations, and an LKJ(1) prior for the residual correlation matrix.

In *Model 2*, we tested the effects of age, sex, aggression initiated and received, dominance rank, and tolerance grade or social style on the expression of SDBs. For dominance rank, we included a penalized cubic regression spline (*k=3*) to capture potential non-linear patterns. An interaction term of rank and grade was used. In the absence of a strong effect, we investigated the individual effects of rank and grade. Although the SDBs were corrected for varying sampling durations, we further checked the effect of observation time. Additionally, we used group size in the model to investigate any potential effects. All three response variables shared the same predictor structure and random-effects structure. We used weakly informative priors for all model parameters. Fixed-effect coefficients were assigned normal (0, 1) priors, and intercepts were given normal priors centred near the empirical means of each response. Residual standard deviations and the standard deviations of all random intercepts were assigned exponential priors with the rate parameter value of 1. The residual correlation matrix among the three SDBs was given an LKJ(2) prior, favoring weakly correlated structures while remaining only mildly informative.

Finally, in *Model 3*, we evaluated the effects of alpha status (alpha vs. non-alpha) on the expression of SDBs, as alpha status may incur an elevated energetic cost compared to intermediate or more subordinate group members [49]. Unlike in other models, here we excluded the data on *M. fuscata* for two reasons: males were not included in the sample, and dominance rank relationships were assessed through matrilineal ranks. Accordingly, we had 12 alpha and 75 non-alpha individuals. Like previous models, we used weakly informative priors for all model parameters. The priors used in this model fully resembled the priors of Model 2.

For sufficient convergence, all Bayesian models were subjected to at least 5500 iterations with 1000 warmup sessions. We reported median estimate coefficients (*Est*), 95% credible interval (*crl*), containing 95% of the posterior probability density function, and probability of direction (*pd*) that indicated certainty of an effect. We established model convergence by following Bayesian statistics guidelines [75]. Furthermore, we checked trace and autocorrelation plots, Gelman-Rubin convergence estimations, and density histograms of posterior distributions. The posterior distributions were sampled using 10,000 iterations and 2000 warmups. We used the “*DHARMa.helpers*” package [76] and investigated model fits by examining the residual distributions (i.e., deviation, dispersion, and outliers). Square-root transformation of yawn was performed for improving model fits of 2 and 3. We applied the Leave-one-out (LOO) cross validation method for model selection based on the expected log pointwise predictive density (ELPD). Only best-fitted models are presented in the results.

## 3. Results

### 3.1. Self-directed behaviours in macaques show only partial correlational consistency

We found only partial correlational consistency among the three SDBs (**Figure 1, Table S2**). Our model, which controlled for species– and group-level structure, revealed a moderate positive residual association between self-scratch and autogroom (*Est* = 0.21, 95% *crl* [0.02 – 0.38], *pd* = 0.98). By contrast, the residual correlations between self-scratch and yawn (*Est* = 0.08, 95% *crl* [-0.12 – 0.27], *pd* = 0.77) and between autogroom and yawn (*Est* = 0.02, 95% *crl* [-0.17 – 0.21], *pd* = 0.57) were weak. Taken together, these results suggest that while some self-directed behaviours may share partially overlapping sources of variation, their associations are generally weak. Thus, the three SDBs do not appear to be driven by a single common underlying factor, supporting our decision to model them as separate response variables rather than combining them into a single composite index.

**Figure 1.**
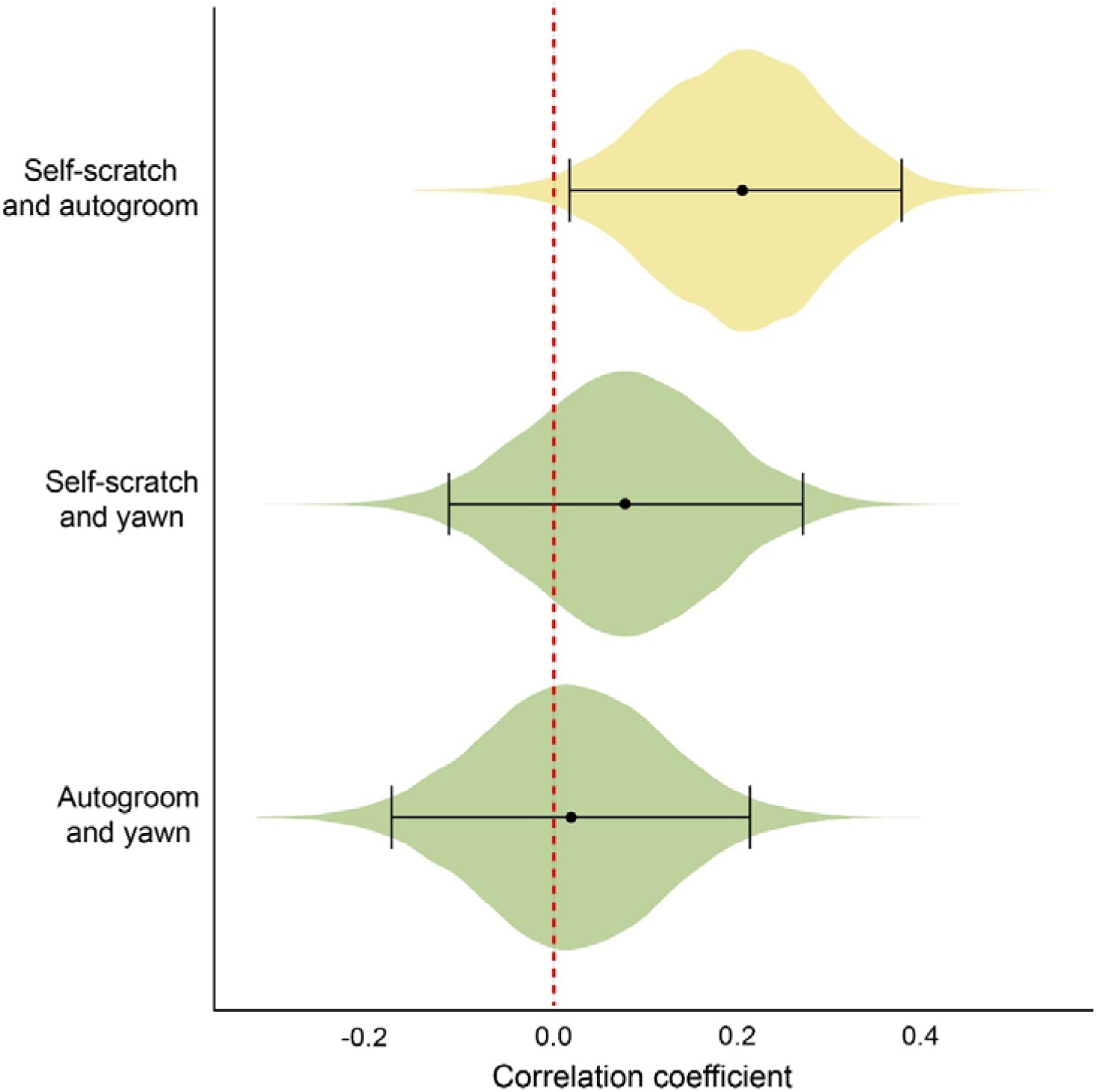
Self-directed behaviours partially correlate with each other. Posterior estimates of residual correlations among self-directed behaviours. Solid black points show the posterior median correlation coefficient for each behavioural pair, while horizontal bars represent the 95% *crl*. The violins show data distribution. The vertical red dashed line indicates a parameter estimate of zero, i.e., the overlap of the *crl* with this line suggests no strong correlations. The yellow color highlights a moderately strong correlation.

### 3.2. Factors associated with self-directed behaviours in macaques

The LOO model comparison identified the multivariate model, which included sex, aggression received, dominance rank, tolerance grade, observation time, and group size, as the best-fitting model for predicting self-scratch, autogroom, and yawn. The dropped variables had no influence on the expression of SDBs. See **Table S3** for model comparison and selection using the LOO method. We found that the key predictors had varying associations with the three self-directed behaviours (**Table S3**).

#### 3.2.1. Associations with sex and aggression

Sex had no effect on self-scratch (*Est* = –0.04, 95% *crl* [-0.08 – 0.01], *pd* = 0.95) and autogroom (*Est* = –0.77, 95% *crl* [-1.82 – 0.30], *pd* = 0.92), but it strongly predicted yawning, with males showing more frequent yawning than females (*Est* = –0.08, 95% *crl* [-0.10 – – 0.05], *pd* = 1.00, **Figure 2**). Aggression received was not associated with self-scratch (*Est* = 0.00, 95% *crl* [-0.03 – 0.03], *pd* = 0.52) and yawn (*Est* = –0.01, 95% *crl* [-0.02 – 0.01], *pd* = 0.83), but was negatively related to autogroom, suggesting that individuals who received lower aggression exhibited prolonged autogroom bouts (*Est* = –0.84, 95% *crl* [-1.61 – 0.09], *pd* = 0.98).

**Figure 2.**
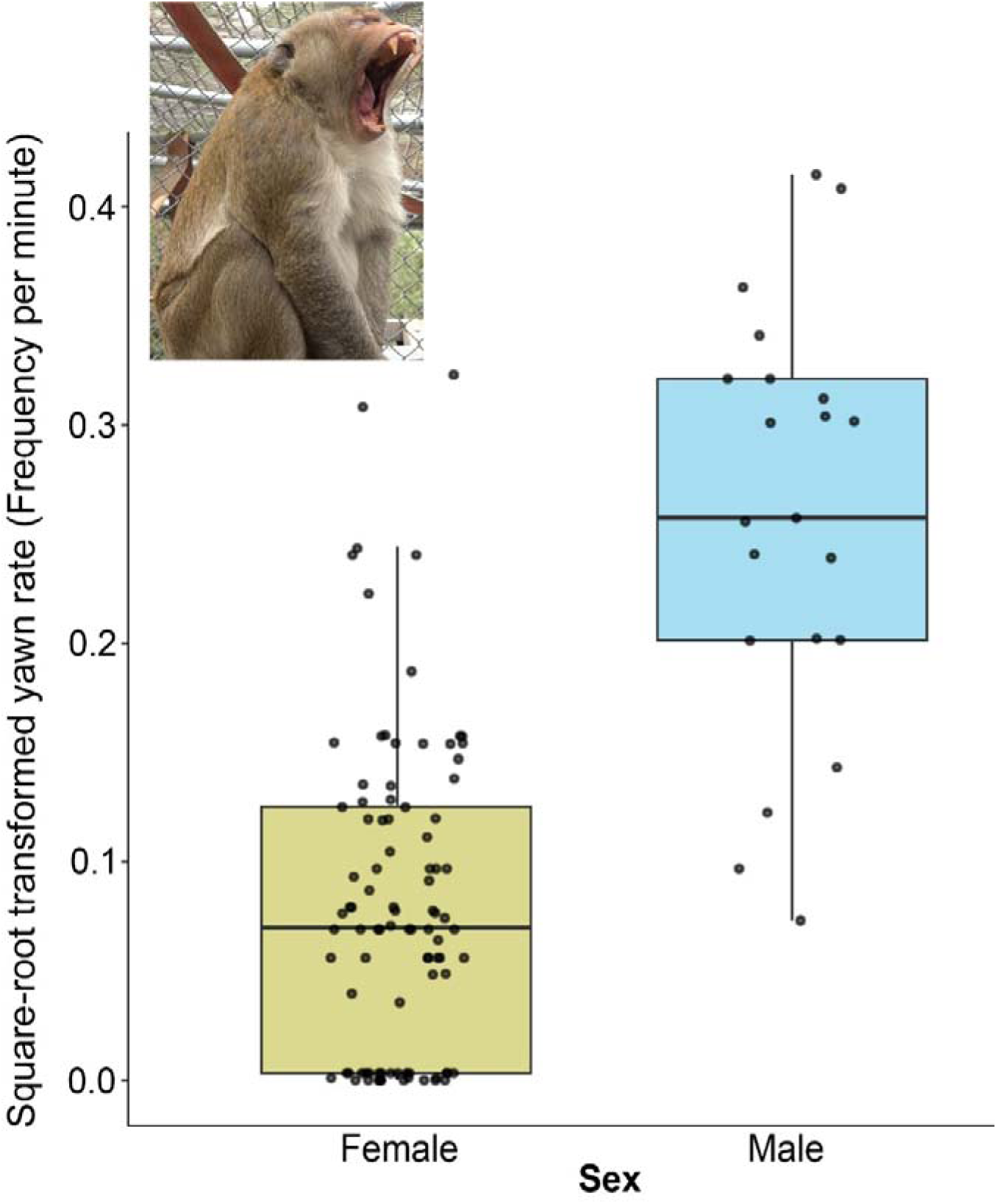
Males exhibit more frequent yawn than females. Square-root transformed yawn rates between male and female macaques. Boxes represent interquartile ranges, and whiskers represent the upper and lower limits of the data. The horizontal bars within the boxes represent the median values. Solid black dots show the distribution of data. Macaque photo credit: Monkey Reality project from Animal Behaviour & Cognition group, Utrecht University.

#### 3.2.2. Associations with dominance rank

We found evidence of both linear and non-linear relationships between dominance rank and the SDBs (**Figure 3**). While our analysis on rank showed limited evidence for an effect on self-scratch (*Est* = 0.03, 95% *crl* [-0.00 – 0.06], *pd* = 0.97, **Figure 3a**), it was strongly associated with autogroom (*Est* = 1.14, 95% *crl* [0.35 – 1.94], *pd* = 0.99, **Figure 3b**) and yawn (*Est* = 0.02, 95% *crl* [0.01 – 0.04], *pd* = 0.99, **Figure 3c**) behaviours. These results suggest that low-ranking individuals engaged in longer autogrooming bouts than their high-ranking counterparts. By contrast, the non-linear relationship between rank and yawning behaviour suggests that yawn rates were lowest at intermediate ranks and increased toward both high– and low-ranking individuals.

**Figure 3.**
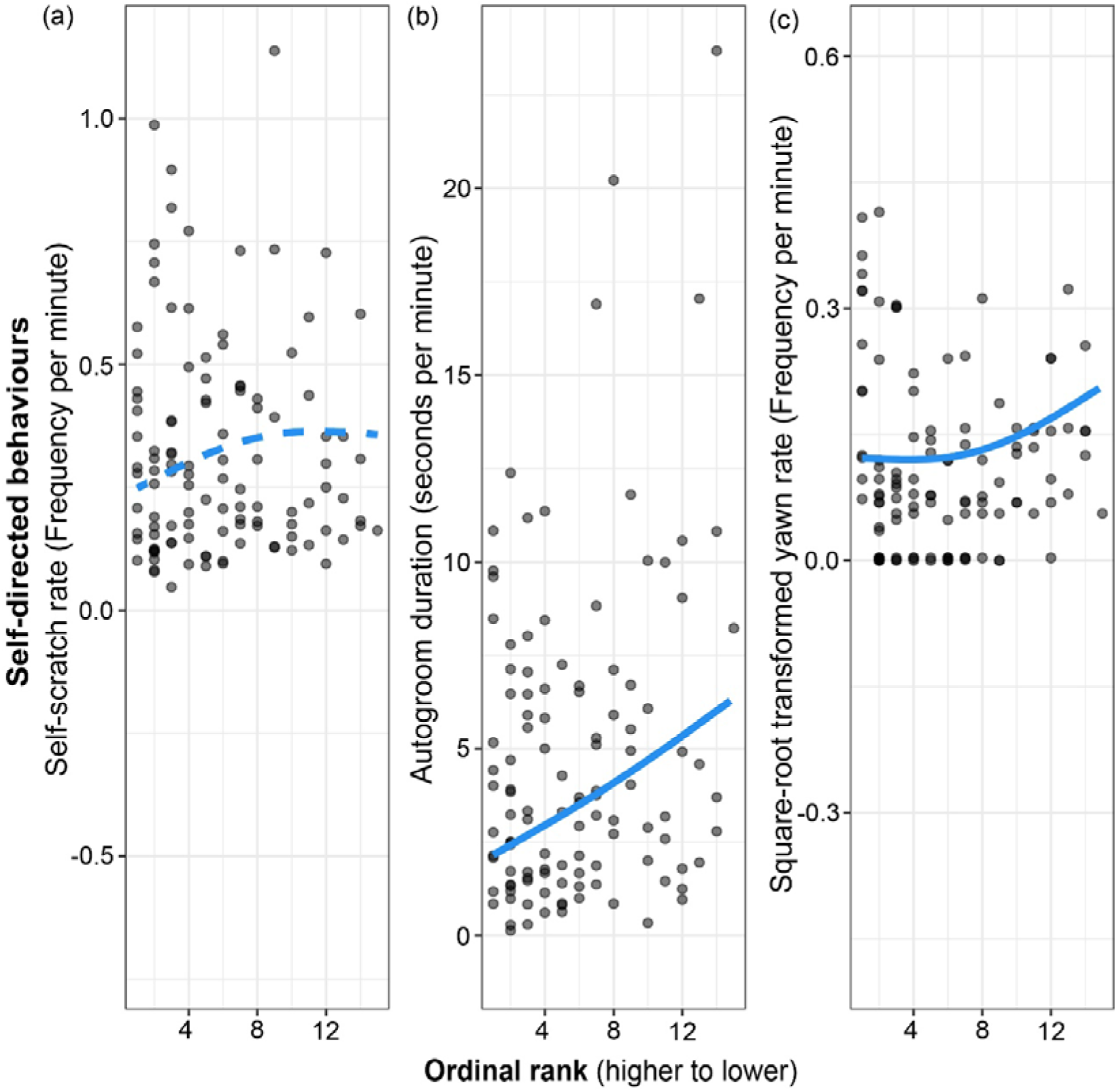
Relationships between dominance rank and self-directed behaviours in macaques. (a) relationship between self-scratch and ordinal rank, (b) relationship between autogroom and ordinal rank, (c) relationship between square-root transformed yawn and ordinal rank. Solid black points show raw data points and overlaid in blue are the posterior predictions. Dashed blue line indicates no strong relationship, whereas the solid lines suggest strong relationship between rank and self-directed behaviours.

#### 3.2.3. Effects of social styles

The mean (±SD) rates of self-scratch were 0.18 ± 0.08, 0.36 ± 0.25, 0.46 ± 0.21, and 0.41 ± 0.22 for Grades 1, 2, 3, and 4, respectively (**Figure 4a**), whereas the mean durations of autogroom bouts were 3.14 ± 2.98, 8.61 ± 5.70, 5.12 ± 3.34, and 3.13 ± 2.52 (**Figure 4b**). Likewise, the mean rates of yawn were 0.01 ± 0.02, 0.02 ± 0.04, 0.03 ± 0.04, and 0.03 ± 0.04 for the four tolerance grades (**Figure 4c**). Despite a visual pattern of the most despotic societies exhibiting lower SDBs, tolerance grade showed no consistent associations, with all estimates highly uncertain and 95% *crl* values spanning zero (**Table S3**). However, we found substantial species– and group-level variation in autogrooming (species SD = 2.17; group SD = 2.12), moderate variation in self-scratch, and minimal variation in yawning (**Table S3**). No strong effects of observation time and group size were found (**Table S3**).

**Figure 4.**
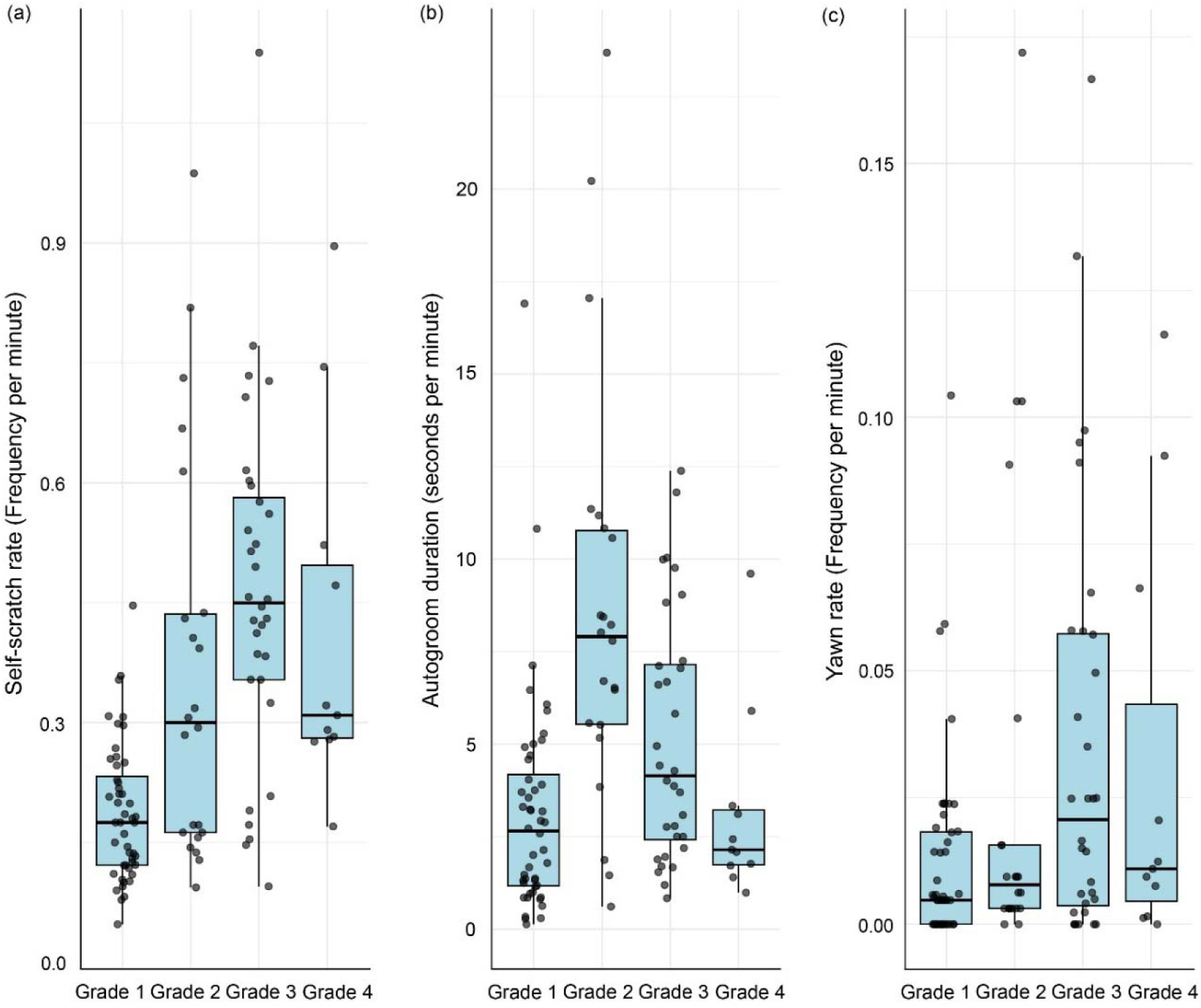
Self-directed behaviours across macaque social styles or tolerance grades. **(a)** rate of self-scratch behaviour, **(b)** duration of autogroom, and **(c)** rate of yawn behaviour. Boxes represent interquartile ranges, and whiskers represent the upper and lower limits of the data. The horizontal bars within the boxes represent median values. Solid black dots show the distribution of data.

### 3.3. Yawn under pressure? Alpha status is linked to increased rates of yawn

The mean (±SD) rates of self-scratch were 0.326 ± 0.154 for alpha individuals and 0.362 ± 0.244 for non-alpha individuals. Similarly, the mean (±SD) durations of autogrooming bouts were 5.105 ± 3.628 for alphas and 4.984 ± 4.533 for non-alphas, while yawning rates were 0.068 ± 0.054 and 0.022 ± 0.032 for alpha and non-alpha individuals, respectively. Alpha individuals did not differ from non-alpha individuals in autogrooming (*Est* = 0.49, 95% *crl* [-1.65 – 2.57], *pd* = 0.63, **Table S4**) or self-scratch (*Est* = –0.06, 95% *crl* [-0.15 – 0.02], *pd* = 0.92, **Table S4**). By contrast, yawning rates were higher in alpha individuals compared to non-alpha (*Est* = 0.12, 95% *crl* [0.07 – 0.17], *pd* = 1.00, **Figure 5**, **Table S4**).

**Figure 5.**
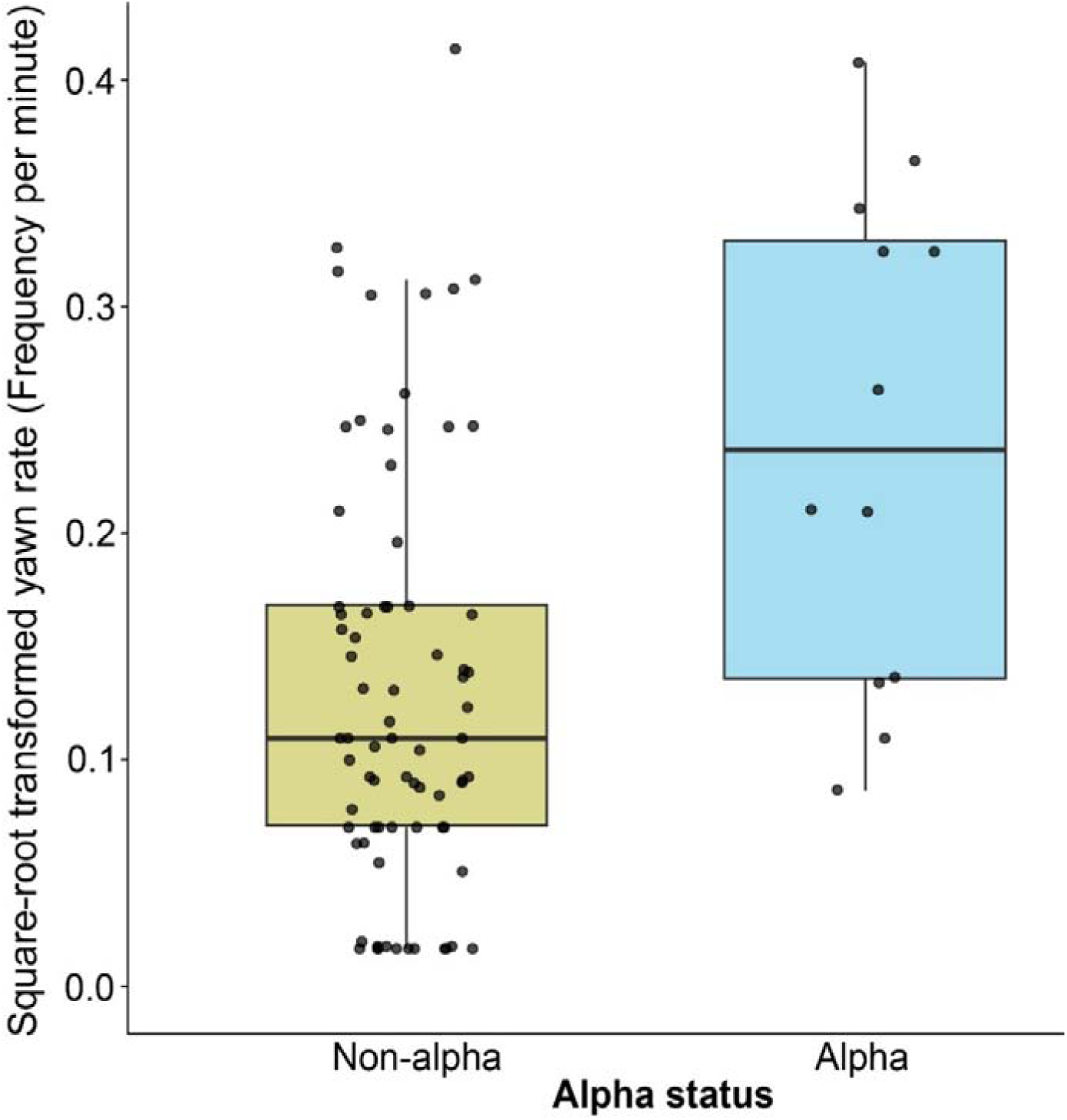
Rate of yawn between alpha and non-alpha individuals. Boxes represent interquartile ranges, and whiskers represent the upper and lower limits of the data. The horizontal bars within the boxes represent the median values. Solid black dots show the distribution of square-root transformed data.

## Discussion

SDBs are considered displacement behaviours that mediate anxiety-like emotional states and serve functions, like assisting individuals to avoid potential conflicts or seek support from closely affiliated group members through visual means [6,13,22,41]. Yet conclusive evidence on the proximate drivers and the associated demographic and social factors underlying variations in SDBs is lacking. Moreover, existing research predominantly includes specific social contexts, like affiliative interactions and conflicts, and often uses social variables like proximity to neighbors (to assess dominance and indirect social influence of group members on others). However, complex emotional states like anxiety are shaped by the continuous and multifaceted processes of social interactions, including broader social dynamics. We conducted behavioural observations on macaques from six different species that varied along a despotic–egalitarian gradient, and investigated three widely recognised SDBs: self-scratch, autogroom, and yawn. Our results suggest that these behaviours share only a limited correlational structure and are influenced differently by demographic and social factors. While we found little to no evidence to support the idea that SDBs are influenced by the broader social dynamics of a species (cf. [50,51]), dominance rank-related effects were complex and more pronounced. These results indicate that SDBs do not form a simple unified anxiety construct and instead may stem from a complex set of functions like social uncertainty, signaling, and hygiene.

### Partially overlapping social functions of self-scratch, autogroom, and yawn

The functions of SDBs can range from social and environmental to hygienic [6–10]. A study comparing the explanatory power of these three hypotheses found stronger support for the parasitic or hygienic model over the other two (cf. [8]). However, they also found support for the prediction that social status (dominance rank) and the uncertainty (presence of nearby group members) of social interactions predict SDBs. In the current study, which investigates the social and emotional aspects of SDBs, we found that not all three behaviours covaried positively, indicating their functional heterogeneity. This is further corroborated by the fact that each of these behaviours was differentially influenced by some of the demographic and social factors we looked at (see later). Consequently, our findings suggest the need for reevaluation of several previous studies that used composite SDB measures as proxies for anxiety, and we strongly advocate for treating the elemental measures separately. Besides, future research should carefully evaluate the context-specific expression of these behaviours.

### Demographic factors are weakly associated with self-directed behaviours

We found no evidence of associations between age and the SDBs. In humans, greater social contact has been found to alleviate anxiety associated with ageing [77]. Thus, the comparable expression of SDBs across the age range of our study animals (4–29 years) potentially indicates that social living conditions may facilitate better emotional regulation [78]. Nonetheless, the lack of association between age and SDBs corroborated the findings of some previous research [79,80]. Unlike age, sex was strongly associated with yawning, but not with self-scratch and autogroom. Males displayed higher rates of yawning than females. A sex-specific bias in yawning can be explained by sexual dimorphism (cf. [33,34]). Since males are more dominant than females in macaque societies, males may exert dominance and signal threats by yawning more, as they possess larger canines than females. Therefore, instead of anxiety, frequent yawning in males may serve as a form of social signaling [69].

### Social factors underlying self-directed behaviours

Contrary to our prediction, we found that individuals who received more frequent aggression from group members engaged in autogrooming for shorter durations. Additionally, there was no association between aggression and self-scratch or yawn. These findings contradict the results of previous research [8,37–39,81]. An inverse relationship between aggression received and autogroom may indicate the presence of alternate coping strategies mediating aggression-related anxiety, like social buffering [47,48]. Furthermore, the non-significant relationships of self-scratch and yawn may indicate that direct social interactions in the form of aggression may not be a prerequisite to elicit anxiety-related emotional states (but see [37–39]), or that these behaviours primarily serve other functions as discussed before.

Social status or dominance rank exhibited complex effects on autogroom and yawn behaviours, consistent with both uncertainty and energetic-constraint hypotheses. The effect of rank on self-scratch was weak but showed an inverse U-shaped non-linear trend. The relatively higher self-scratch rates among intermediate-ranking individuals could potentially be due to the high probabilities of rank mobility, whereas the highest and lowest ends of the spectrum remain somewhat reserved [82]. By contrast, the prolonged autogrooming bouts in subordinate individuals clearly indicates an influence of social uncertainty (but see [81]). Finally, a U-shaped relationship pattern emerged between rank and yawn, which could be attributed to both rank-related uncertainty (i.e., for lower ranks) and the energetic costs (i.e., for higher ranks) of social ranks (cf. [38,41,44]). The elevated rates of yawning among higher-ranking individuals indicate the heightened physiological demands of mating, vigilance, and group maintenance. This was even further strongly supported by the finding of increased yawning among alpha individuals in comparison to the non-alpha (cf. [49]). In particular, frequent yawning by alpha individuals can convey social signals of dominance to other group members [83,84], which can help maintain the stability of social groups. Alternatively, although not mutually exclusive, yawns are known to function as a brain cooling mechanism and may become more frequent in alpha individuals due to their high loads of vigilance [85]. While elevated rates of yawning in higher-ranking individuals can reflect on certain energy demands, it might be indicative of different emotional costs in lower-ranking individuals. Thus, it is possible that the lower-ranking individuals are probably ‘certain that they are uncertain’ in predicting the outcomes of social interactions (also see [86]), subsequently leading to elevated rates of yawning. These results may also suggest the probable mechanisms of vigilance and social tension at play for subordinate individuals. Nonetheless, the functions of yawning are multifaceted, and more interdisciplinary research is required to assess its role regarding social dynamics. In summary, the rank-related effects and their directions were not consistent for self-scratch, autogroom, and yawn, further stressing the differential mechanisms governing these behaviours in macaques.

Our results revealed no strong social style level variations in self-scratch, autogroom, and yawn. However, individuals belonging to the most despotic social style (i.e., Grade 1) showed a tendency to exhibit lower levels of these behaviours than others (cf. Figure 4). Nonetheless, these patterns are contradictory to the prediction that despotism inflicts pressure on individuals in the form of power asymmetries and frequent aggressive interactions, and that individuals are highly susceptible to negative emotions like anxiety [52,54,59,87]. In line with our findings, a comparative study between despotic long-tailed and egalitarian Tonkean macaques (*Macaca tonkeana*) showed that long-term cortisol concentrations (used as an indicator of stress) were higher among individuals of Tonkean than those of long-tailed macaques [61]. While harsh leadership (such as in despotic societies) can induce stress among humans [88], macaques belonging to the despotic social styles can rely on alternative mechanisms to alleviate and cope with anxiety, such as forming and maintaining selectively strong bonds [89–91] as a social buffer [92] (*sensu* social buffering [47]) and engaging in discriminate cooperative and prosocial interactions [60,93–97]. By contrast, in egalitarian societies, such inter-individual bonds are not necessarily predictable, strong, and stable [59–61,98], potentially dampening the positive effects of tolerance [48]. Moreover, a recent study using the same groups (except the Japanese macaques) as the current, experimentally showed that co-feeding tolerance levels were higher in more despotic societies (cf. [60]). Therefore, the strong selective bonds in despotic societies might have been translated into higher tolerance, which in turn resulted in less frequent exhibition of SDBs.

The alternative prediction, i.e., higher social uncertainty in more egalitarian than despotic societies, was partially supported, at least based on the same trend in the most despotic society. Due to the predictable nature of social relationships in despotic societies, individuals are certain about the possible outcomes of social interactions, subsequently exhibiting lower levels of SDBs. By contrast, higher uncertainty in less despotic or more egalitarian societies can lead to individuals showing greater levels of SDBs. Notably, social uncertainty is considered a critical measure of social complexity [64,65]. Thus, in other words, we can assume that social complexity may play a role in driving anxiety-like behaviours, which should be tested in future studies.

## Ethics

We received ethical approval from the Animal Experiments Committee and the Animal Welfare Organisation at the Biomedical Primate Research Centre (BPRC) (approval no. 019E). At each participating zoo, internal ethical committees further approved and carefully monitored the research procedures. Observations were conducted from a considerable distance, without deliberately influencing the behaviour of the animals. The entire study was strictly non-invasive in accordance with European Directive 2010/63, and we followed the guidelines set forth by the American Society of Primatologists regarding the appropriate treatment and inclusion of animals.

## Data accessibility

Upon publication, data and code will be made publicly available.

## Declaration of AI use

We have not used AI-assisted technologies in creating this article.

## Authors’ contributions

D.B.: conceptualization, methodology, investigation, formal analysis, writing – original draft, writing – review and editing, visualization, supervision, project administration, funding acquisition; E.S.J.vD.: investigation; R.E.M.dV.: investigation; C.E.K.: investigation; M.V.: investigation; A.N.Z.: investigation; J.J.M.M.: conceptualization, methodology, writing – original draft, writing – review and editing, supervision, project administration, funding acquisition. All authors gave final approval for publication and agreed to be held accountable for the work performed therein.

## Acknowledgements

We thank William Allen, Elena Belli, Paula Escriche Chova, Jolanda de Jong, Aníta Guðjónsdóttir, Elja Jeunink, Penny Kuijer, Esmee Middelburg, Veera Schroderus, Sjoerd Sijbrandij, Paola Meems, Amber Kozanli, Yesper Bos, and Mary Maximiadi for their assistance with data collection and organization. We thank the staff members of the Affenberg Zoobetriebsgesellschaft mbH (Austria), Biomedical Primate Research Centre (the Netherlands), Diergaarde Blijdorp (the Netherlands), Gaia Zoo (the Netherlands), Apenheul Primate Park (the Netherlands), Planckendael Zoo (Belgium), and Artis Zoo (the Netherlands) for their support during our research. We thank Peter Gaubatz, Svenja Gaubatz, Jan Langermans, Annet Louise Louwerse, Jos Hartog, Linda Bruins-van Sonsbeek, Emile Prins, Lisette van den Berg, Marjolein Osieck, Lena Pflüger, Karline Janmaat, and Elisabeth Sterck for facilitating this research at respective zoos and institutions. We also thank George Hodgson and Alan McElligott for providing valuable comments on an earlier version of the manuscript.

## Electronic supplementary information to

**Table S1.** Summary of study species and groups with sampling and demographic information.

| Species | Group ID | Focal sampled | Age (years) | Sex |
| --- | --- | --- | --- | --- |
| *Macaca fuscata*<br>(Japanese) | Affenberg | N = 26; 280<br>min/focal | 4 – 29 (median<br>age = 10) | Female = 26;<br>Male = 0 |
| *Macaca mulatta*<br>(Rhesus) | R3G2 | N = 12; 160<br>min/focal | 4 – 14 (median<br>age = 5) | Female = 11;<br>Male = 1 |
|  | R3G7 | N = 10; 210<br>min/focal | 4 – 25 (median<br>age = 13) | Female = 9;<br>Male = 1 |
| *Macaca fascicularis*<br>(Long-tailed) | J1G4 | N = 8; 320<br>min/focal | 4 – 9 (median<br>age = 4) | Female = 7;<br>Male = 1 |
|  | J1G7 | N = 10; 320<br>min/focal | 4 – 14 (median<br>age = 12.5) | Female = 9;<br>Male = 1 |
|  | NWR | N = 4; 320<br>min/focal | 4 – 6 (median<br>age = 4) | Female = 0;<br>Male = 4 |
| *Macaca silenus*<br>(Lion-tailed) | Blijdorp | N = 3; 240<br>min/focal | 6 – 18 (median<br>age = 7) | Female = 2;<br>Male = 1 |
|  | Apenheul | N = 6; 340<br>min/focal | 7 – 27 (median<br>age = 18.5) | Female = 5;<br>Male = 1 |
| *Macaca sylvanus*<br>(Barbary) | Gaia | N = 14; 240<br>min/focal | 4 – 20 (median<br>age = 10.5) | Female = 8;<br>Male = 6 |
|  | Apenheul | N = 9; 420<br>min/focal | 4 – 20 (median<br>age = 7) | Female = 8;<br>Male = 1 |
| *Macaca nigra*<br>(Crested) | Blijdorp | N = 3; 460<br>min/focal | 5 – 21 (median<br>age = 13) | Female = 2;<br>Male = 1 |
|  | Artis | N = 3; 320<br>min/focal | 9 – 21 (median<br>age = 15) | Female = 3;<br>Male = 0 |
|  | Planckendael | N = 5; 180<br>min/focal | 4 – 15 (median<br>age = 6) | Female = 2;<br>Male = 3 |

The *M. fuscata* group was located at Affenberg Landskron Park in Villach, Austria, within a large 40,000 m² enclosure characterized by native mixed woodlands typical of Southern Austria. This group comprised approximately 170 individuals. Two *M. mulatta* groups (R3G2 and R3G7), with 30 and 25 members respectively, were housed at the BPRC in Rijswijk, the Netherlands. Both groups lived in facilities featuring identical enclosure sizes: 74 m² indoors and 250 m² outdoors. At the same facility (BPRC), we also studied three *M. fascicularis* groups: J1G4 (15 individuals), J1G7 (18 individuals), and NWR (4 individuals). J1G4 and J1G7 had access to the same enclosure sizes (49 m² indoor and 183 m² outdoor), while the smaller NWR group was housed in smaller enclosures measuring 3.55 m² indoors and 3.88 m² outdoors. The two groups of *M. Silenus* were housed at Blijdorp Zoo and Apenheul Primate Park in the Netherlands. The Blijdorp group comprised 3 individuals and had 100 m² indoor and 106 m² outdoor spaces, whereas the Apenheul group included 8 individuals, with access to an 80 m² indoor enclosure and a significantly larger 768 m² outdoor area. Two *M. sylvanus* groups were included: one at Gaia Zoo in the Netherlands and the other at Apenheul Primate Park. The Gaia Zoo group consisted of 14 individuals and had access to a large 3522 m² outdoor area and a 108 m² indoor space. The Apenheul group, with 13 individuals, lived exclusively in a 3829 m² outdoor enclosure featuring natural elements such as vegetation, rocks, and a creek. Finally, the three *M. nigra* groups were housed at Blijdorp Zoo, Artis Zoo (both in the Netherlands), and Planckendael Zoo in Belgium. Group sizes were 5, 4, and 6 individuals, respectively. The Blijdorp group was kept in a 70 m² indoor and a 160 m² outdoor enclosure. At Artis Zoo, the enclosures measured 65 m² indoors and 761 m² outdoors, while the Planckendael group had access only to an 82 m² indoor enclosure.

In all cases where both indoor and outdoor spaces were provided, macaques could freely move between them via connecting tunnels. An exception was the Apenheul *M. sylvanus* group, where only the outdoor space was available during observations. Group sizes include all members, including infants under one year of age.

Although husbandry practices varied slightly between groups—reflecting differences in species-specific needs and institutional protocols—all followed the standards set by the European Association of Zoos and Aquaria for animal housing and care. Enclosures were equipped with various enrichment items, including climbing structures, ropes, tree trunks, wooden elements, and slides. With the exception of the *M. fuscata* and Apenheul *M. sylvanus* groups, which were kept exclusively in outdoor habitats, indoor housing areas were climate-controlled and featured concrete floors layered with sawdust bedding. Feeding schedules differed across groups to meet specific dietary requirements but remained consistent with their usual routines during our observations. Standard diets included commercial monkey pellets, a variety of fruits and vegetables, and seed mixes such as sunflower and corn.

**Table S2.**
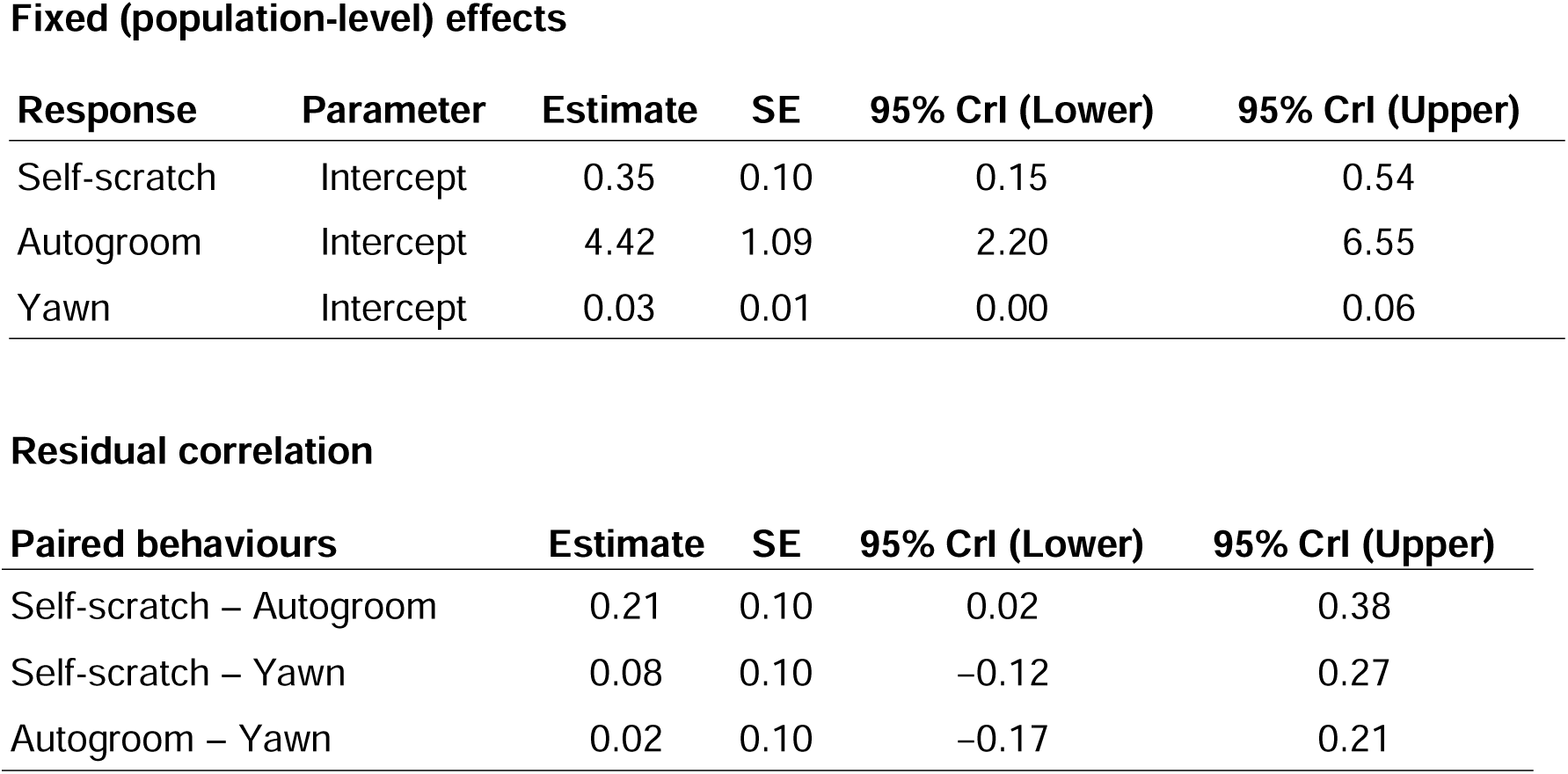
Summary of Bayesian multivariate hierarchical model (*Model 1*) investigating the correlational consistency among self-scratch, autogroom, and yawn behaviours.

**Table S3.** Summary of Bayesian multivariate hierarchical model (*Model 2*) investigating the effects of sex, aggression received, dominance rank, and tolerance grade or social style on self-scratch, autogroom, and yawn behaviours.

**Self-scratch**
| Predictor | Estimate | SE | 95% CrI (Lower) | 95% CrI (Upper) |
| --- | --- | --- | --- | --- |
| Intercept | 0.31 | 0.36 | -0.66 | 0.92 |
| Grade 1 | 0.04 | 0.80 | -1.12 | 2.25 |
| Grade 2 | 0.01 | 0.75 | -1.71 | 1.50 |
| Grade 3 | -0.04 | 0.62 | -1.56 | 1.03 |
| Aggression received | 0.00 | 0.02 | -0.03 | 0.03 |
| Sex (male) | -0.04 | 0.02 | -0.08 | 0.01 |
| Mean observation | 0.02 | 0.08 | -0.15 | 0.18 |
| Group size | -0.24 | 0.61 | -2.05 | 0.54 |

**Autogroom**
| Predictor | Estimate | SE | 95% CrI (Lower) | 95% CrI (Upper) |
| --- | --- | --- | --- | --- |
| Intercept | 3.46 | 1.31 | 0.64 | 5.73 |
| Grade 1 | -4.08 | 3.08 | -10.72 | 1.75 |
| Grade 2 | 3.17 | 2.74 | -2.76 | 8.65 |
| Grade 3 | 0.70 | 2.32 | -3.82 | 5.65 |
| Aggression received | -0.84 | 0.39 | -1.61 | -0.09 |
| Sex (male) | -0.77 | 0.53 | -1.82 | 0.30 |
| Mean observation | -0.29 | 0.83 | -1.89 | 1.42 |
| Group size | 0.82 | 2.22 | -3.44 | 5.67 |

**Yawn**
| Predictor | Estimate | SE | 95% CrI (Lower) | 95% CrI (Upper) |
| --- | --- | --- | --- | --- |
| Intercept | 0.13 | 0.23 | -0.47 | 0.53 |
| Grade 1 | 0.08 | 0.44 | -0.63 | 1.19 |
| Grade 2 | -0.03 | 0.46 | -1.07 | 0.89 |
| Grade 3 | -0.01 | 0.37 | -0.83 | 0.67 |
| Aggression received | -0.01 | 0.01 | -0.02 | 0.01 |
| Sex (male) | -0.08 | 0.01 | -0.10 | -0.05 |
| Mean observation | -0.02 | 0.03 | -0.07 | 0.03 |
| Group size | -0.13 | 0.25 | -0.83 | 0.19 |

**Smooth terms (dominance rank)**
| Response | Smooth term | Estimate | SE | 95% CrI (Lower) | 95% CrI (Upper) |
| --- | --- | --- | --- | --- | --- |
| Self-scratch | s(ord_rank) | 0.03 | 0.02 | -0.00 | 0.06 |
| Autogroom | s(ord_rank) | 1.14 | 0.40 | 0.35 | 1.94 |
| Yawn (sqrt) | s(ord_rank) | 0.02 | 0.01 | 0.01 | 0.04 |

*Note:* Best-fitted model presented (Model3). Model selection was done using LOO method.
Result of LOO comparison of models –
elpd\_diff se\_diff
Model5 0.0 0.0
Model4 -0.3 0.5
Model3 -0.6 1.1
Model2 -3.0 2.2
Model1 -3.9 2.8
Variables included in Models:
**Model1:** s(ord\_rank, k = 3) + grade + agginitiate + aggreceive + age + sex + meanobs + groupsize + (1 | species / group)

**Table S4.** Summary of Bayesian multivariate hierarchical model (*Model 3*) investigating the alpha vs non-alpha individuals on self-scratch, autogroom, and yawn behaviours.

| Predictor | Estimate | SE | 95% CrI (Lower) | 95% CrI (Upper) |
| --- | --- | --- | --- | --- |
| Intercept | 0.38 | 0.13 | 0.13 | 0.62 |
| Alpha vs non-alpha | -0.06 | 0.05 | -0.15 | 0.02 |

**Autogroom**
| Predictor | Estimate | SE | 95% CrI (Lower) | 95% CrI (Upper) |
| --- | --- | --- | --- | --- |
| Intercept | 4.40 | 1.26 | 1.81 | 6.89 |
| Alpha vs non-alpha | 0.49 | 1.07 | -1.65 | 2.57 |

**Yawn**
| Predictor | Estimate | SE | 95% CrI (Lower) | 95% CrI (Upper) |
| --- | --- | --- | --- | --- |
| Intercept | 0.12 | 0.04 | 0.05 | 0.19 |
| Alpha vs non-alpha | 0.12 | 0.03 | 0.07 | 0.17 |

